# Chromosome-level assembly of the Arabian killifish as a novel biomedical model species

**DOI:** 10.64898/2026.09.15.751479

**Authors:** Ruth Y. Akinmusola, Tetsuo Kon, Koto Kon-Nanjo, Rashid Minhas, Paul O’Neill, Aaron Jeffries, Yasuhito Shimada, Mark Ramsdale, Tetsuhiro Kudoh

## Abstract

The Arabian killifish (*Aphaniops dispar*) is an excellent model for studying human diseases such as fungal infections and cancer progression. The embryos possess a transparent chorion for live imaging and display an extended period of independent feeding (13 days post-fertilisation). They also exhibit broad thermal tolerance which enables live imaging at physiologically relevant human temperatures. However, omics resources for this species remain limited and restricts its use in genomics studies. We generated a 1.45 GB high-quality reference assembly with 24 chromosome-level scaffolds and N50 score of 60.64 Mb by combining long-read, long-range, and short-read sequencing. The *A. dispar* genome exhibits a high level of homozygosity (95.6%) with BUSCO gene completeness scores of 99.2% and 97.5% for the Actinopterygii and Cyprinodontiformes lineages, respectively. The genome is highly repetitive (58.7%) and the majority of these are DNA transposons. The mitochondrial genome is 81.55% similar to that of *Aphanius iberus* (the Spanish toothcarp), with minor structural modifications. *A. dispar* exhibits highly conserved synteny with *Fundulus heteroclitus* within the Cyprinodontoidei clade. Together, these findings provide a valuable genomic resource for comparative genomics within the teleost community.

## Introduction

Teleost species within the Cyprinodontoidei lineage exhibit remarkable ecological resilience, which enables them to thrive in dynamic and extreme environments^1^. The Arabian killifish (*Aphaniops dispar*; synonym: *Aphanius dispar*) is a relatively small sized Cyprinodontoid widely distributed in ecosystems with fluctuating temperatures and broad salinity gradients^2,3^. The species have been found in ecosystems ranging from fresh water desert ponds to hypersaline coastal lagoons across the red sea, Arabian Peninsula and Persian Gulf^2,4^. These habitats can experience rapid and substantial environmental shifts such as flash floods which promote rapid physiological and molecular acclimation in fish^2^. Additionally, adult *A. dispar* exhibit pronounced thermotolerance, maintaining normal gonadal development and reproductive activity across a broad temperature range (18°C and 37°C) without evidence of reproductive impairment^5^. Interestingly, the embryos remain viable at high temperatures (37°C and 42°C) for limited periods. The tolerance to environmental stressors positions *A. dispar* as a model for manipulating temperature, salinity and oxygen levels to study responses to ecological changes and host-pathogen interactions^6^. This offers a unique advantage to study immune or physiological responses at human-relevant and fever-range temperatures, a limitation to traditional teleost models like Zebrafish^7^. Unlike Zebrafish, *Aphaniops dispar* exhibits slower embryo and larval development, resulting in an extended period prior to the onset of independent feeding at 13 days post fertilisation^5^. The prolonged developmental window offers potential refinement and reduction benefits for conducting experimental work before the animals reach a protected feeding stage under animal-welfare regulations.

The *A. dispar* evolutionary history suggests its suitability as a model for studying climate-driven diversification and immune gene evolution. Phylogeographic evidence revealed substantial intraspecific differentiation over time among populations from different geographic locations including the Persian Gulf and the Gulf of Oman^8^. Environmental changes such as sea level fluctuations and geographic fragmentation can cause shifts in separated and reconnected *A. dispar* populations^9^. This ecological plasticity and the ability to adapt to local environment conditions provides a useful way to compare population responses to changes such as temperature variation, salinity shifts or pathogen exposure^2,10^. The embryos are also suitable models for environmental risk assessment with high sensitivity to a wide range of contaminants such as surfactants and disinfectants^11^. Recent work has shown that *A. dispar* embryos display transcriptional and physiological responses in response to pathogenic infection and environmental changes within hours to days^2,6,11^. Additionally, *A. dispar* populations from contrasting salinity regimes show differences in osmoregulatory, ion transport and immune-related gene expression^2^. Furthermore, these pathways are dynamically regulated during salinity transitions indicating a broad adaptive framework for coping with hyperosmotic stress^2^. This makes the species a great model for studying climate change related disease susceptibility or immune-gene evolution across divergent populations.

Despite the progress in establishing *A. dispar* as a promising experimental model, key gaps have slowed its advancement compared to other killifish models. Well established killifish species such as *Fundulus heteroclitus, Nothobranchius furzeri* and *Kryptolebias marmoratus* possess high-quality reference genomes, population-level resequencing datasets and extensive functional genomics tools^12–16^. In contrast, *A. dispar* lacks a chromosomally complete reference genome and comprehensive genomics resources restricting its use in evolutionary analysis and environmental adaptation studies. This limits genotype-to-phenotype mapping experiments, selection scans and the detection of adaptive loci. To overcome these limitations, we generated the first chromosome-scale reference genome for *A. dispar*. This approach integrated Nanopore long read, Illumina short read and Omni-C chromatin conformation data. The assembly spans 24 chromosomes, consistent with the expected karyotype for Aphaniidae members^17,18^. The resultant genome shows high gene completeness establishing *A. dispar* as a genomics-enabled model species. The availability of this reference genome provides a foundation for deeper functional studies, including immune-related gene evolution, identification of stress-response pathways and population-level analysis on lineage-specific diversification of adaptive immune functions.

## Methodology

### Fish preparation for DNA sequencing

The animal procedures in this study were carried out according to the UK Home Office relevant guidelines and regulations with the licence number PP4402521. All the procedures were conducted with respect to ARRIVE guidelines and appropriate measures taken to reduce pain, suffering and distress to the animals throughout the study. An eighteen-month-old Arabian killifish was reared in captivity at the Aquatic Resources Centre, the University of Exeter. Breeding conditions were controlled at a constant temperature of 28 °C, and a light cycle of 14 h of light, subsequently 10 h of darkness. Culling experiments were carried out under schedule1 procedure for termination of small fish using termination by benzocaine overdose in system water and confirmation of death with brain destruction.

### Library construction and whole-genome sequencing

All DNA sequencing libraries were derived from a single eighteen-month-old Arabian killifish male. For both Illumina and Nanopore sequencing, muscle tissue was dissected from the male sample, immediately flash frozen in liquid Nitrogen before storage at −80 °C. High molecular weight (HMW) DNA was extracted using the Monarch HMW DNA Extraction kit for tissue (New England Biolabs, catalogue number NEB #T3010). The integrity and concentration of the DNA was analysed using the Agilent fragment analyser and Qubit dsDNA system. Short reads up to 25Kb were eliminated using the Circulomics short read eliminator kit prior Nanopore sequencing. The ONT ligation sequencing kit was used for library construction using the size selected genomic DNA. Long read sequencing was performed using the Nanopore PromethION PCA100193 flow cell in the sequencing facility at the University of Exeter. Evaluation of the resultant Nanopore reads with QUAST v5.0.2^19^ revealed 4389 contigs with a N50 of 3.26Mb and a total GC content of 39.52% (Table S1). Short read genomic sequencing was performed with the Illumina Novaseq6000 sequencer and S1 flowcell (2 x 150bp). The quality of the short reads was checked with the FastQC v0.12.1^20^ and summarised with multiQC v1.35^21^. Low quality and adapter sequences were removed from the reads with fastp v0.23.4 with default parameters which retained 237.4 million paired end reads with GC content of 39% and a total of 70.4 Gb.

### Omni-C library construction, sequencing and scaffolding

Approximately 100 mg of fish muscle was ground to fine powder in liquid nitrogen using a mortar and pestle. The ground tissue was sequentially crosslinked with disuccinimidyl glutarate and formaldehyde in PBS. After washing, chromatin was digested with the DNase-based Nuclease Enzyme Mix supplied with the Dovetail Omni-C Kit (Cantata Bio, CA, USA). The digested chromatin was captured on Chromatin Capture Beads and subjected to end polishing, bridge ligation, and intra-aggregate ligation. Crosslinks were then reversed and the DNA was purified. Illumina-compatible libraries were subsequently prepared according to the manufacturer’s protocol. The Omni-C library was sequenced on the Illumina NovaSeq X Plus sequencing platform, generating a total of 42,275,382 paired-end Omni-C reads. Read quality was assessed using FastQC v0.12.1.

For scaffolding, the Omni-C reads were aligned to the draft genome and filtered for valid contacts using Juicer v1.6^22^ with the -s none option. Scaffolding was performed using 3D-DNA v180419^23^ with option “-r 2”. The resultant scaffolds were manually curated with Juicebox Assembly Tools v1.11.08^24^. The final assembly size is 1.45 Gb containing 24 chromosome-scale scaffolds with a N50 of 60.64Mb and an overall anchor rate of 98.4% (Figure 1, Table S2). This is consistent with earlier karyotyping studies, which reported 24 chromosomes in the Arabian killifish^17,25^. The scaffolds were then polished with Illumina genomic short-reads using Pilon v1.24^26^. Finally, the assembly was screened for cobionts and symbionts using NCBI FCS-GX^27^ and all sequences identified as contaminants were masked.

**Figure 1.**
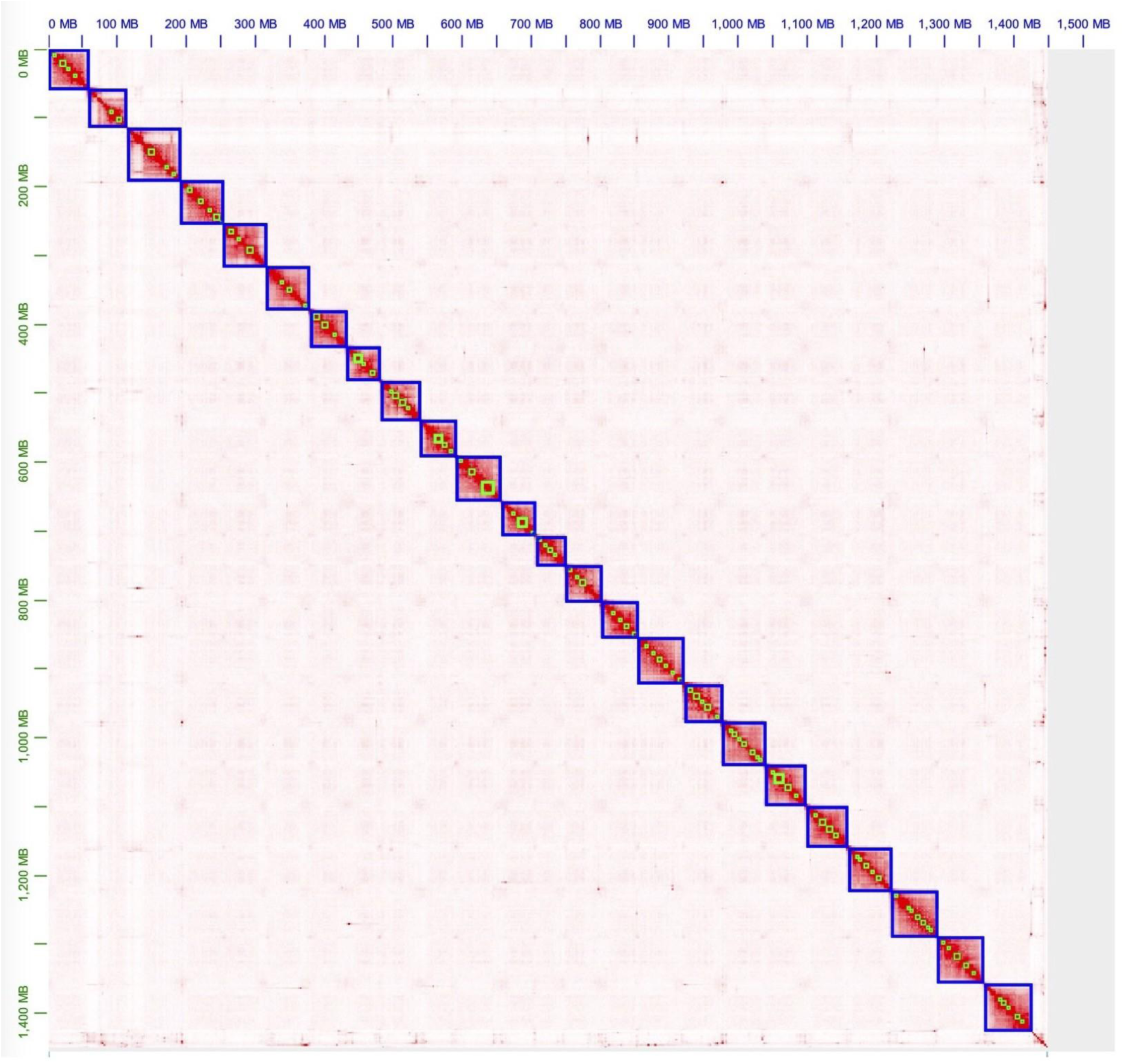
Hi-C contact map for the chromosome-level genome assembly of the Arabian killifish. The red spots indicate chromatin contact signals and their intensity revealing the strength of chromatin contact. The total number of chromosome-scale scaffolds is 24.

### DNA metabarcoding and mitogenome extraction

We analysed all the contigs assembled for the *A. dispar* genome using the protocol described for the doctor fish^28^. BLASTN v2.16.0^29^ was used to screen for sequences homologous to the *Aphanius iberius* mitochondrial sequence query (OP884090.1, length=16,708bp) from the NCBI database^30^. We identified a single BLAST hit from contig_8643 with a total length of 16521bp (Table S3). The hit aligned throughout the entire length of the *A. iberius* mitogenome with total sequence identity of 82% and 1% gap. We annotated this contig with the MitoAnnotator web server^31^ which revealed 13 protein coding genes, 2 ribosomal RNA genes (12s and 16s), 22 transfer RNA genes and a D-loop (Figure 2). This mitogenome aligns well with the mitochondrial genomes of the Cyprinodontoidei close relatives on the NCBI with the highest sequence identity of 99% with that of Cyprinodon *variegatus* (Table S4, Figure S1). Insertions and deletions are the major differences distinguishing the mtDNA of *A. dispar* from that of other Aphaniidae members (Table S4). Mitogenome phylogenetic analysis placed the Arabian killifish within the Aphaniidae genus in the Cyprinodontoidei lineage. The strongest relationship was observed with *Aphanius iberus* within the Aphaniidae clade (Figure 3 and S1).

**Figure 2.**
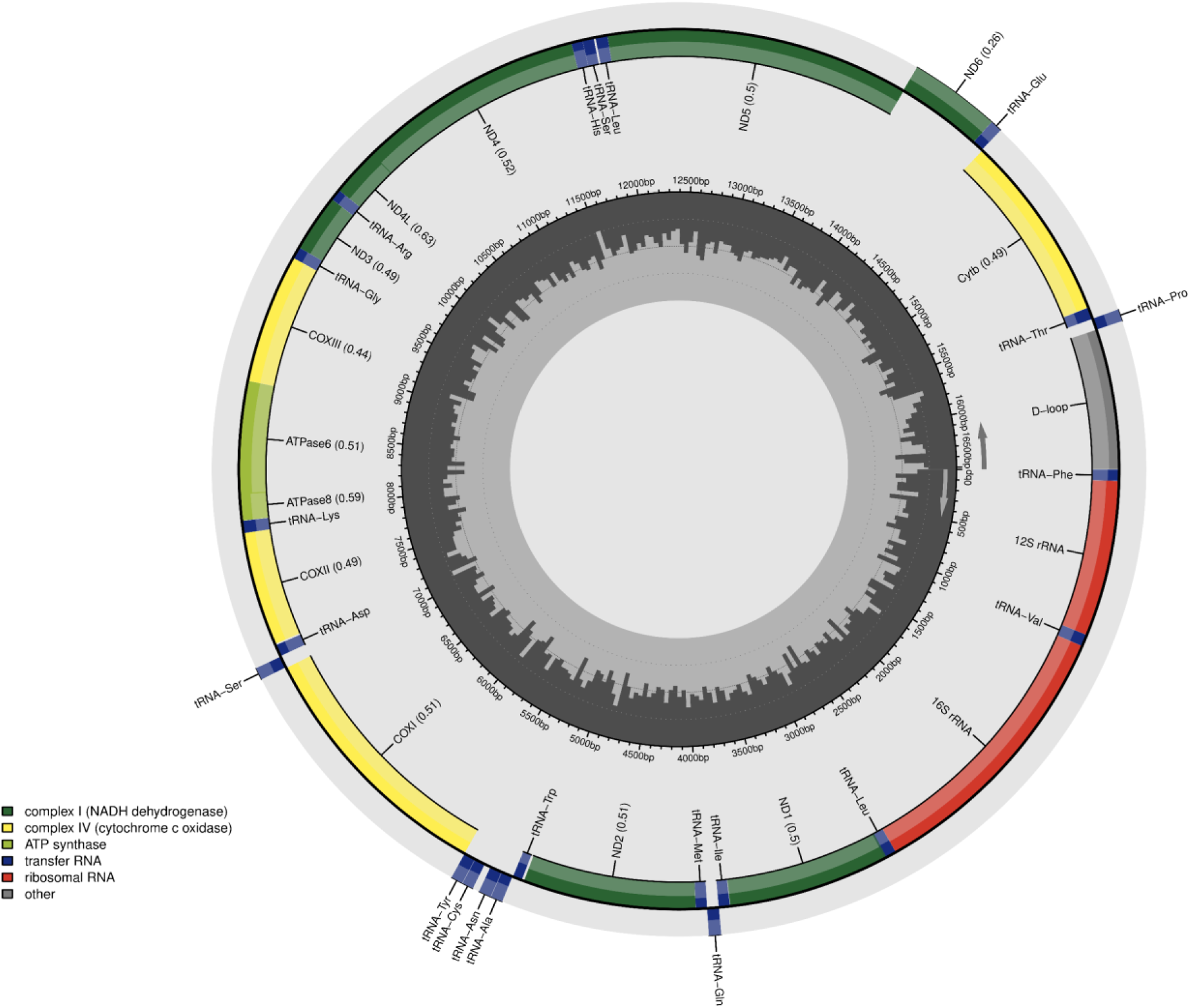
*Aphaniops dispar* Mitochondrial genome. A circular representation of the mitochondrial genome of *A. dispar.* It contains 13 protein-coding genes, 2 ribosomal RNA genes (12s and 16s), 22 transfer RNA genes and a D-loop. Starting from the outermost circle, all the protein-coding genes are depicted on the scale in deep green, light green and yellow colours. The D-loop is in grey, the 2rRNA genes are illustrated in red, while the 22tRNA genes are depicted in blue illustrations. The inner circle with black colour shows the length in base pairs (bp).

**Figure 3.**
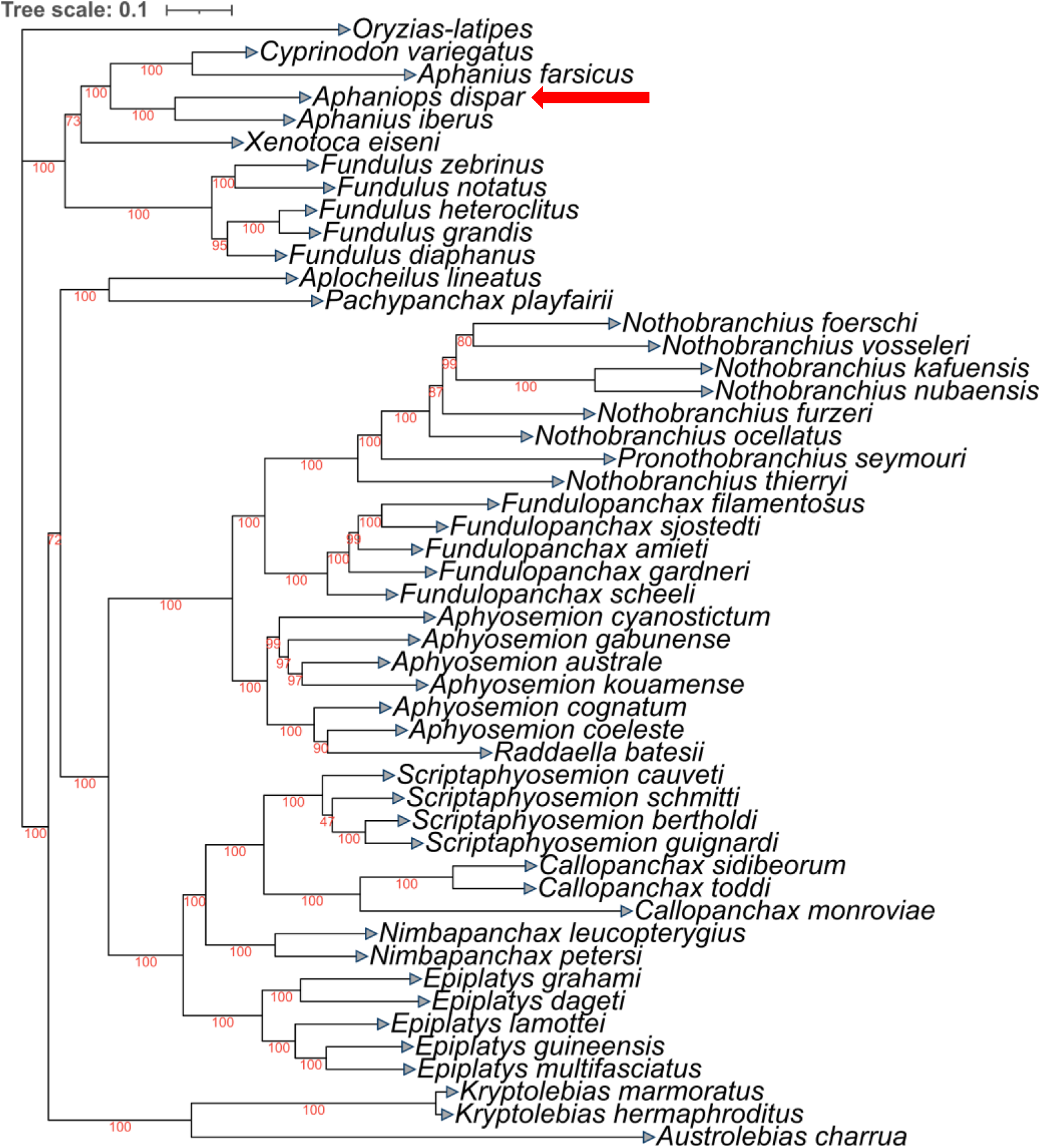
A maximum likelihood tree revealing the phylogeny of the *A. dispar* mitogenome within different killifish species. Bootstrap support values on the branches are indicated in red. The scale bar indicates an average phylogenetic distance of 0.1 substitutions per site. The red bar is associated with *A. dispar* mitogenome position within the phylogenetic tree.

#### Genome size estimation

The *A. dispar* genome size was estimated using the k-mer profiles of the Illumina reads analysed with Jellyfish 2.3.1^32^. Computation of genome size and heterozygosity was carried out using genomescope v2.0^33^ based on the 21-mers estimated using Jellyfish. GenomeScope was used to estimate the genome size, heterozygosity, ploidy level and error rate from the computed 21-mers. The genome is diploid with an estimated genome length of 1.04Gb (Figure 4). The genome is estimated to be homozygous at most loci with very low heterozygosity (Figure 4). The 95.6% homozygosity is close to the expectations from a highly inbred fish but below the threshold reported for medaka inbred lines^34^.

**Figure 4.**
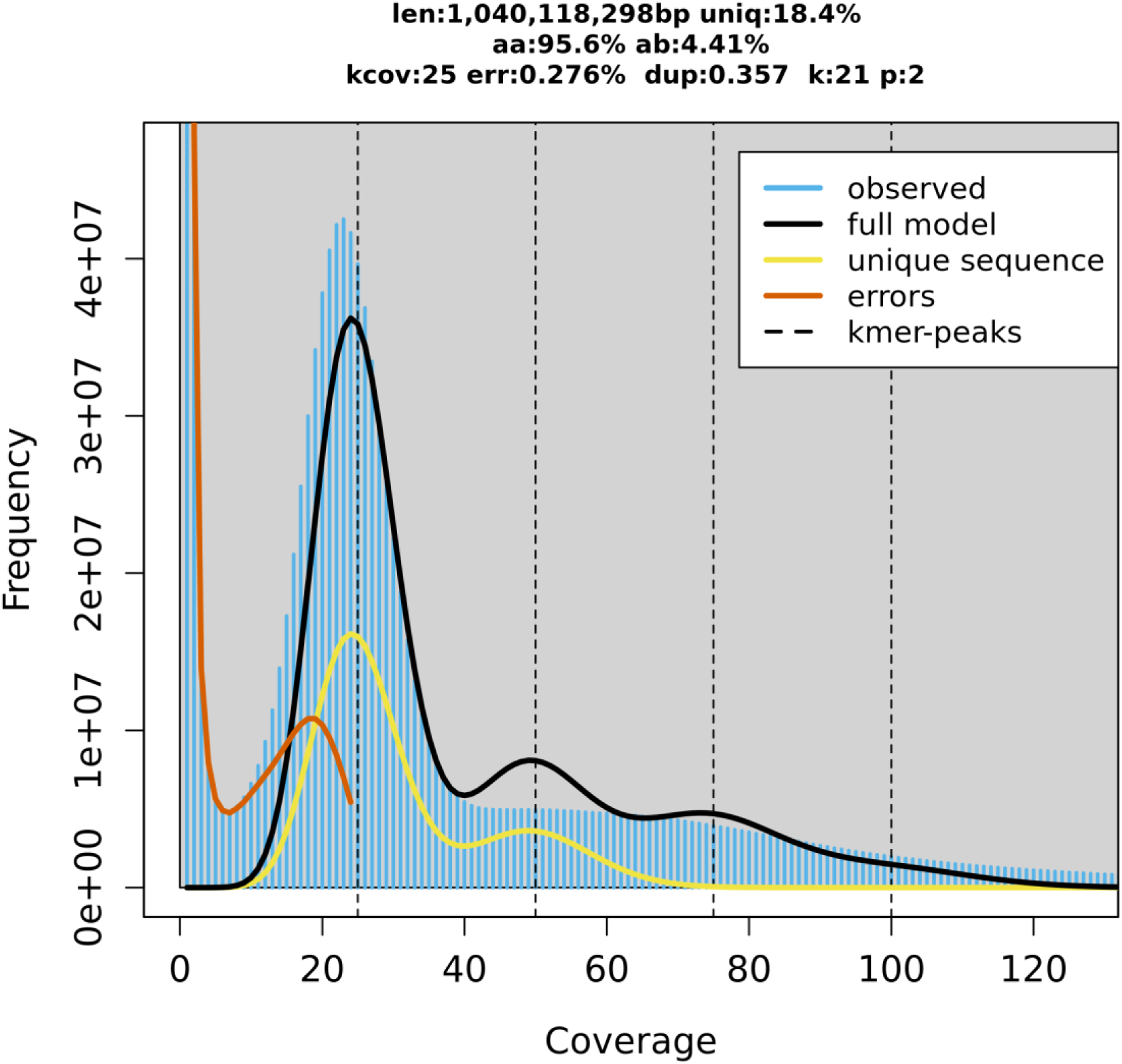
The k-mer profiles of the Illumina short reads based on k value of 21. The x-axis represents the k-mer coverage and the y-axis represents the frequency of k-mers per coverage. Len: The estimated genome size is 1.04Gb. Uniq: The frequency of unique alleles is 18.4%. AA: The estimated homozygosity is 95.6%. AB: The estimated heterozygosity is 4.41%. kcov: the estimated coverage of unique genomic k-mers. Err: error rate of 0.276%. dup: duplication levels of 0.357. k: 21 k-mers. P: ploidy level is diploid.

### Repeat annotation

Repeat elements were annotated with the EarlGrey repeat annotation pipeline version 4.4.4^35^ within a conda environment. EarlGrey used RepeatModeler v2.0.5^36^ to generate a non-redundant repeat library and the resultant library was fed to RepeatMasker v4.1.5^37^ for repeat masking. Repbase partitions were used during the Repeatmodeler and Repeatmasker integration steps to reorganise TE predictions for accurate classifications and non-overlapping genomic partitions. LTR_FINDER was employed under the EarlGrey implementation for full-length and intact LTR retrotransposon identification. The *A. dispar* genome is repeat-rich (58.7%), mainly dominated by DNA transposons (21.78%), LINE elements (13.26%). A total of 17.53% unclassified repeats were accounted for (Figure 5A, Table 1).

**Figure 5.**
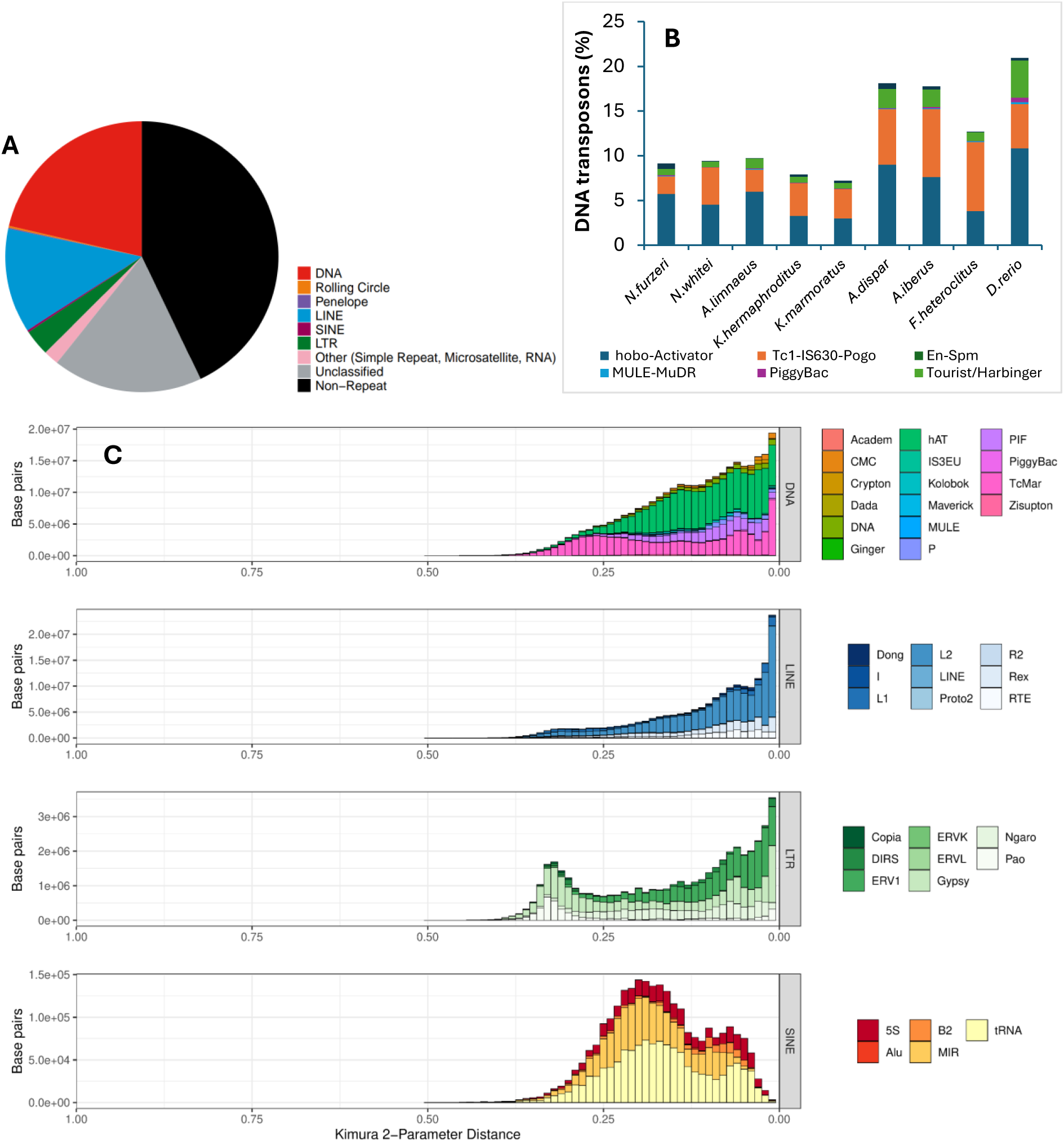
Repeat content and landscape in the *A.dispar* assembly. (**A**) The proportion of repeats in the entire assembly. (**B**) The distribution of different DNA transpons within the entire assembly compared with the EarGrey annotation output of some fish genomes. (**C**) The landscape of repetitive elements within the chromosome-scale scaffolds. The x-axis represents the Kimura parameter distance, which estimates the evolutionary divergence of the younger repeat elements from their consensus sequences. The y-axis is the relative proportion of genome (in base pairs) occupied by each diverged repetitive element.

**Table 1.** The proportion of repetitive elements in the *A. dispar* assembly.

|  | All scaffolds |  |  | Chromosomes |  |  | Unplaced sequences |  |  |
| --- | --- | --- | --- | --- | --- | --- | --- | --- | --- |
| Repetitive element type | Number of elements | Length (bp) | Genome coverage (%) | Number of elements | Length (bp) | Genome coverage (%) | Number of elements | Length (bp) | Genome coverage (%) |
| DNA | 994,717 | 315,705,644 | 21.78 | 943,870 | 305,887,620 | 21.45 | 6,204 | 2,521,656 | 10.69 |
| Rolling circle | 8,083 | 7,643,663 | 0.53 | 9,239 | 6,137,640 | 0.43 | 712 | 1,543,231 | 6.54 |
| Penelope | 3,713 | 1,456,409 | 0.10 | 5,914 | 1,988,958 | 0.14 | 0.00 | 0.00 | 0.00 |
| LINE | 420,762 | 192,192,560 | 13.26 | 411,673 | 188,222,573 | 13.20 | 6,316 | 2,755,030 | 11.68 |
| SINE | 17,481 | 3,438,871 | 0.24 | 16,871 | 2,852,718 | 0.20 | 234 | 156,187 | 0.66 |
| LTR | 86,583 | 48,194,185 | 3.33 | 89,806 | 48,754,318 | 3.42 | 847 | 424,043 | 1.798 |
| Other* | 353,982 | 28,249,404 | 1.95 | 347,028 | 25,922,134 | 1.81 | 942 | 4,035,010 | 17.11 |
| Unclassified | 753,226 | 254,017,167 | 17.53 | 780,187 | 249,865,006 | 17.52 | 6,836 | 3,592,990 | 15.24 |

\*Other (Simple Repeat, Microsatellite, RNA)

Most of these repetitive elements were successfully anchored into their chromosomal locations with the complete anchorage of all Penelope elements (Table 1). The DNA repeat signature in the *A. dispar* genome is similar to that of *Aphanius iberus* and some other teleosts with the enrichment of Tc1-IS630-Pogo and hobo-Actvator elements (Figure 5B). The repeat landscape of the Arabian killifish revealed a recent burst of young transposons which were detected in the entire assembly (Figure S5A), chromosome-level scaffolds (Figure 5C) and unplaced scaffolds (Figure S5B). In the chromosome-level scaffolds, the majority of these young transposons are DNA, LINE and LTR transposons (Figure 5C). A relatively higher proportion of diverged yonger transposons elements were detected in the chromosome-level scaffolds when compared to the unplaced scaffolds (Figure 5C and S5B).

Additionally, the chromosomal sequences were screened for telomeric repeats using quarTeT^38^ TeloExplorer version 1.2.4 with these flags: -c animal (TTAGGG repeat unit), and -m 100 and -m 25 (long and truncated telomeric repeats). The telomeric repeats identified were reported on the reverse complement strand as AACCCT, which represents a rotational variant of the C-rich vertebrate telomere repeat (TTAGGG/CCCTAA). Only one chromosome (Chr 15) contained long telomeric repeat arrays at both ends (≥600bp), with seven others (1, 2, 6, 10, 11, 19 and 24) having these repeats at one end (Table S5-S6). We also detected long telomeric repeat arrays at the ends of five unplaced scaffolds but their relative sizes are within the 1Mb range suggesting these as collapsed telomeric repeat artifacts or misjoined chromosome ends. Twenty-two of the chromosomes have at least a quarter (≥150bp) of the AACCCT telomeric repeats at both ends with the exception of chromosomes 18 and 23 (Table S7).

### Gene annotation and functional prediction

Gene prediction was carried out with BRAKER3 using transcriptome and protein evidence. The transcriptome evidence was derived from Illumina RNA-seq reads of the four embryonic stages (11dpf, 7dpf, 5dpf and 72hpf) highly aligned to the softmasked *A. dispar* genome with HISAT2 v2.2.1^39^ with default settings (Table 2, Table S9). The protein sequences from 26 Cyprinodontiformes and Cyprinodontoidei members were used for homology prediction (Table S8). Braker3 was run in a conda environment integrating GENEMARK-ETP and AUGUSTUS prediction with the zebrafish training set. Compleasm was incorporated directly into the BRAKER3 pipeline using the Cyprinodontiformes_odb10 BUSCO lineage flag to assess annotation completeness. Final gene models were generated by TSEBRA, which reconciled predictions into a high confidence consensus annotation. Corresponding gff3 files was generated from the BRAKER3 gtf output with genometools. We predicted a total of 24,360 protein coding genes in the genome and annotation statistics were computed with AGAT (Another GFF analysis Toolkit) v1.4.1^40^ using the longest isoforms (Table S10). The predicted genes models exhibit an average coding sequence length of 1.67 kb, average gene length of 24.05 kb and a mean exon length of 174 bp (Table S10).

**Table 2.** Annotation metrics and proteome consistency of the BRAKER3 annotated *A. dispar* genome.

| **BRAKER3 Annotation metrics** |  |
| --- | --- |
| RNA-seq evidence | 11 dpf, 7 dpf, 5 dpf and 72 hpf embryonic transcriptome |
| Protein evidence | 26 Cyprinodontiformes and Cyprinodontoidei members |
| BRAKER3 --cyprinodontiformes flag? | Yes |
| **BUSCOs for longest isoforms** |  |
| BUSCO (actinopterygii_odb10) | C:99.2%[S:98.6%,D:0.6%],F:0.2%,M:0.6%,n:3640 |
| BUSCO (cyprinodontiformes_odb10) | C:97.5%[S:96.7%,D:0.8%],F:0.5%,M:2.0%,n:15213 |
| **Transcript statistics** |  |
| Mono:Multi exon Ratio | 0.1 |
| Number of genes | 24, 360 |
| Number of exons per transcript | Minimum: 1<br>25%: 4<br>50%: 8<br>75%: 14<br>Maximum: 260 |
| **Omark metrics (longest isoforms)** |  |
| Single | 17,012 (92.2%) |
| Duplicated | 520 (2.82%) |
| Duplicated, unexpected | 509 (2.76%) |
| Duplicated, expected | 11 (0.06%) |
| Omark missing HOGs | 920 (4.99%) |
| Consistent lineage placements | 21,992 (90.30%) |
| Inconsistent lineage placements | 1109 (4.55%) |
| Omark known proteins | 23,101 (94.85%) |
| Omark unknown proteins | 1,254 (5.15%) |
| Total Contaminants | 0.00% |
| Clade detected | Cyprinodontoidei |
| Number of conserved HOGs | 18,452 |

BUSCO: Benchmarking Universal Single-Copy Orthologs, C: Complete BUSCOs, S: Complete and single-copy BUSCOs, D: Complete and duplicated BUSCOs, F: Fragmented BUSCOs, M: Missing BUSCOs, n: Total BUSCO groups searched, dpf: days post fertilisation and hpf: hours post fertilisation. For the Omark proteome consistency, genes in the unknown do not share enough similarity with known gene families within the Cyprinodontoidei clade. These could include orphan genes or erroneous annotations.

The peptide sequences from the annotation were blasted (BLASTP, BLASTX) against the Cyprinodontiformes and Cyprinodontoidei members used for annotation and the UniProtswissprot database (Figure 6A, Table S12-S16). Furthermore, eggNOG-mapper v2.1.12^41^ and Interproscan^42^ were used to functionally annotate the BRAKER3-predicted protein sequences (Figure 6B, Table S17-S18). The maximum overall functional annotation rate of 99.6% was achieved in the BLASTP search against the UNIPROT-SwissProt database followed by 99.47 annotation rate from BLASTX search against the 26 Cyprinodontiformes RefSeq genomes and the least annotation rate of 96.06% from the eggNOG database (Figure 6B). Consequently, BLASTP search against the UNIPROT-SwissProt database and BLASTX search against the 26 Cyprinodontiformes RefSeq genomes exhibited the highest unique annotation rate of 55 and 15 additional hits respectively (Figure 6B). The overall annotation was summarised with BLASTP hits in the UNIPROT-SwissProt database mapped to UniProtKb annotation, GProfiler Zebrafish orthologues and Zebrafish blast hits with respect to their chromosomal location and gene ontology terms (Table S19).

**Figure 6.**
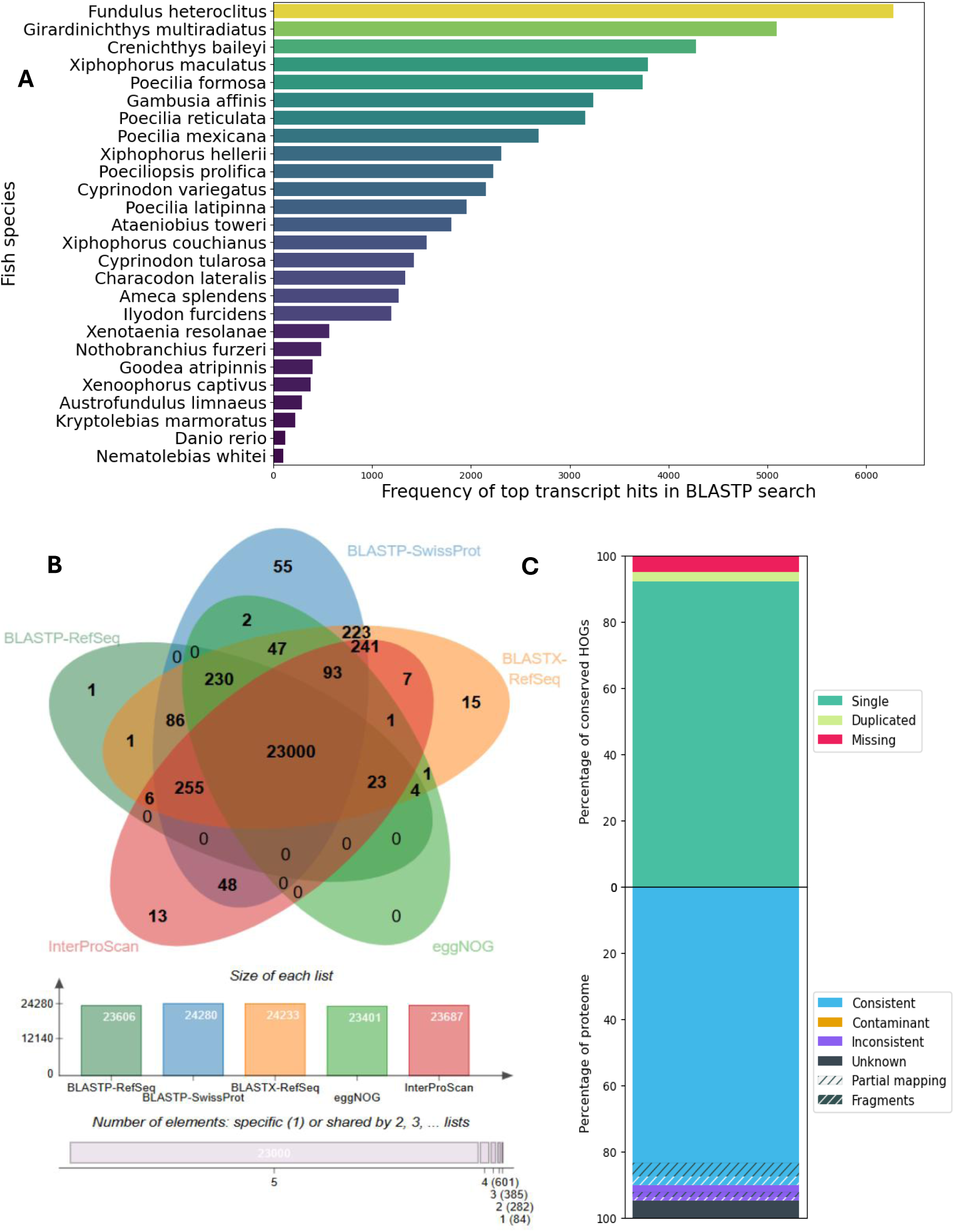
Gene functional annotation statistics and taxonomy profiles. (**A**) The bar plot shows the relative frequency of BLASTP hits against different fish proteomes. (**B**) The Venn diagram shows functional annotation statistics with BLAST, eggNOG Mapper and InterProScan. (**C**) Omark proteome consistency assessment for all the annotated proteins based on Hierarchical Orthologous Groups (HOGs) within the Cyprinodontoidei clade.

We combined Omark v0.4.1^43^ and BUSCO v5.8.3^44^ analysis for assessing annotation gene completeness. Proteome consistency of the resultant annotation was analysed with Omark which detected the Cyprinodontoidei taxonomic clade with no contaminant sequences. The results on the 18,452 conserved Hierarchical Orthologous Groups (HOGs) revealed 17,012 (92.20%) single copies, 520 (2.82%) duplicated copies, 509 (2.76%) duplicated but unexpected, 11 (0.06%) duplicated and expected copies, and 920 (4.99%) missing copies (Figure 6C and Table 2). The annotated assembly possessed a BUSCO gene completeness of 99.2% for the Actinopterygii_0db10 lineage with negligible missing BUSCOs (0.5%) among the total 3,640 BUSCO groups searched (Table 2 and S11).

### Orthologous cluster identification and phylogeny

The proteomes of nine teleost species were downloaded from the NCBI FTP genome datasets. These species include *Cyprinodon variegatus*, *Gambusia affinis*, *Xiphophorus maculatus*, *Girardinichthys multiradiatus*, *Poecilia reticulata*, *Poecilia formosa*, *Crenichthys baileyi* and *Danio rerio* as an outgroup (Table S20). Single copy orthologues and gene family clusters were identified using the Orthofinder algorithm^45^ in an all against all BLASTP searches. The single copy orthologue sequences were extracted from orthofinder and independently aligned using MAFFT v7.505^46^. The alignment was trimmed to remove poor regions with trimAL v1.4.1^47^ and concatenated into a supermatrix alignment plus partitions file with AMAS tool^48^ for downstream phylogenetic analysis. Phylogenetic inference was performed with IQ-TREE v2.2.2.3^47^ with partitioned model finder and branch support assessed with ultrafast bootstrap replicates. The maximum likelihood method was implemented in IQTREE and the resultant tree file was visualised using iTOL (the Interactive Tree Of Life) v7.6^49^ with leaf sorting applied to improve visual clarity. This placed *A. dispar* within the Cyprinodontoidei clade with a monophyletic relationship to *Gambusia affinis. Xiphophorus maculatus* and the live-bearing Poecilids (Figure 7A).

**Figure 7.**
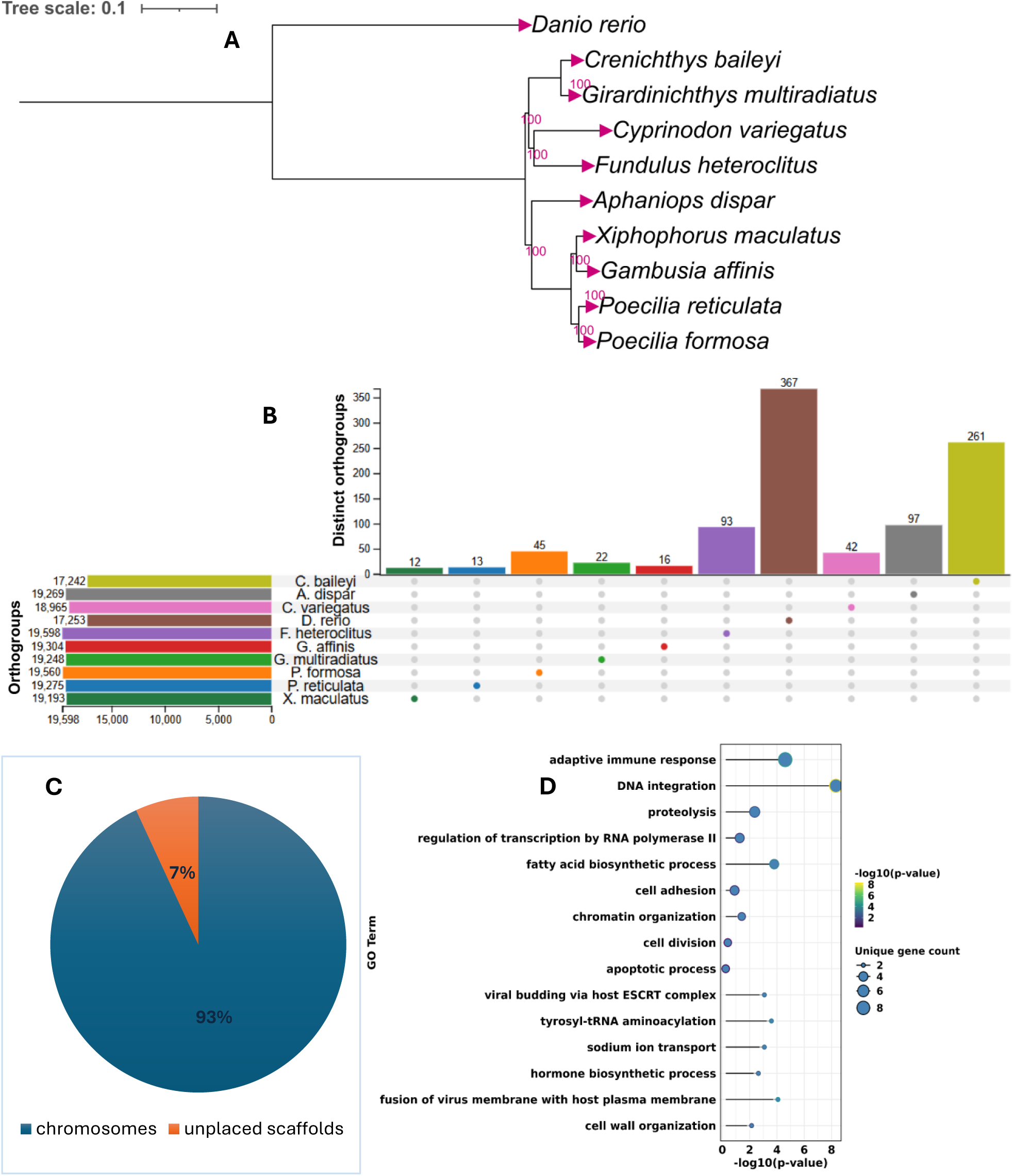
Comparative overview of orthologue clusters from all annotated protein sequences in the Arabian killifish genome. A. Maximum likelihood phylogenetic tree inferred from a concatenated alignment of single copy orthologue sequences using IQ-TREE. B. An upset plot showing the total number of orthologroups and distinct orthogroups in the Arabian killifish and nine other teleost genomes. C. The distribution of *A. dispar* specific orthogroups in chromosomes and unplaced sequences. D. GO annotation showing the top significantly enriched biological processes within the *A. dispar* specific orthogroups.

Furthermore, we identified 97 *A. dispar* specific orthologues with GO terms from disease-relevant pathways and mostly non-zebrafish annotations (Figure 7B, S6, and Table S21-S24). Interestingly, the majority of these are from chromosomal scaffolds (Figure 7C). They are involved in biological processes such as WNT signalling pathway, response to oxidative stress, regulation of immune response, and adaptive immune response. The adaptive immune response cluster is among the top significantly enriched biological processes (Figure 7D). Top hits in this GO term are human orthologues of T cell receptor loci (TVAL3-HUMAN) with no hits in the RefSeq fish genomes used for annotation (Table S19 and S24). Nested within these TCR loci are immunoglobulin orthologues and a small number of them are within the top significantly enriched cellular processes (extracellular region) in Figure S7 and Table S25.

Some of these TCR members also belong to some other GO terms and their lack of characterisation in other fish genomes could signify highly variable or rapidly evolving TCR loci within the Arabian killifish genome. A typical T-cell receptor sequence consists of many segments (VDJC) and the variations in these segments are driven by recombination, not sequence conservation^50,51^. This is because the rearranged sequence is not inherited in the next generation^50^. While human T cell receptor loci have been extremely well curated^52^, the corresponding teleost homologs remain under-annotated and are frequently designated as uncharacterised proteins (Table S19 and S24). Thus, the enrichment observed here, reflect the diversification of the fish adaptive immune response rather than contamination or annotation artifacts. This may indicate selective pressures associated with pathogen exposure or ecological niche.

### Chromosome synteny analysis reveals extensive conservation between *A. dispar* and representative Cyprinodontoidei genomes

Chromosomal homology using collinear blocks for all possible chromosome pairs were computed with the MCScanX algorithm using whole-genome reciprocal BLASTP results with evalue 1e-5 and max_target_seqs 1^53^. Collinear gene sets comparing *A. dispar* with *Danio rerio*, *Fundulus heteroclitus* and *Girardinichthys multiradiatus* were extracted with MCScanX. The corresponding collinear results and gff3 coordinates were visualised with synvisio^54^. The syntenic blocks between *A. dispar* and the cyprinodontoidei members (*F. heteroclitus* and *G. multiradiatus*) are more conserved with lesser fragments than those observed with *D. rerio* (Figure 8). The reduced synteny observed with *D. rerio* may reflect the evolutionary divergence between Cypriniformes (represented by the Cyprinid *D. rerio*) and Cyprinodontiformes (the order containing many killifish species)^55^. Together, these chromosome-scale comparisons validate the quality of the *A. dispar* genome assembly and demonstrate a broad conservation consistent with teleost genomes while highlighting clade-specific structural divergence.

**Figure 8.**
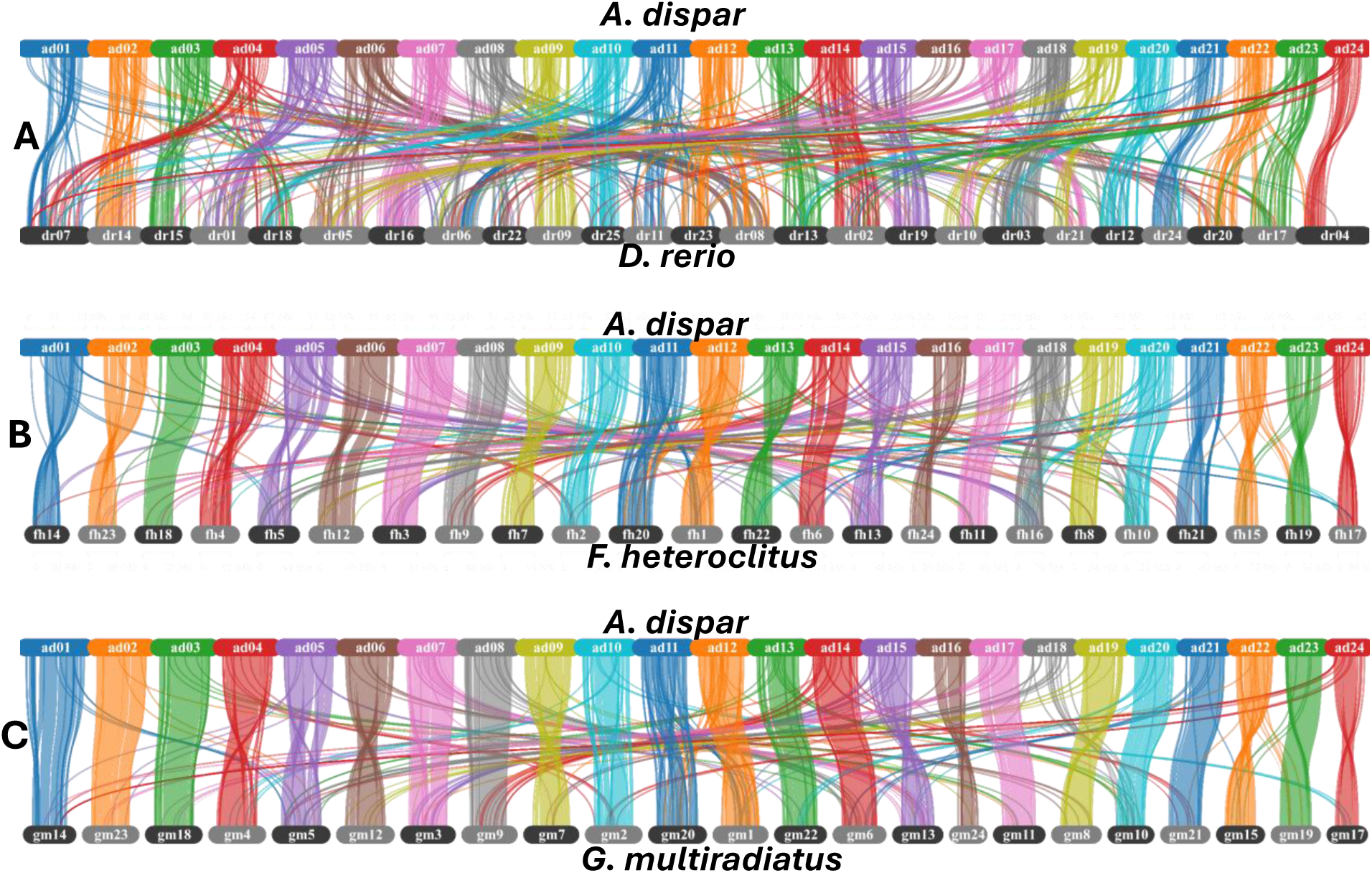
Syntenic relationships between A. dispar and representative teleost species. A. *A. dispar* vs *D. rerio*. B. *A. dispar* vs *Fundulus heteroclitus* C. *A. dispar* vs *G. multiradiatus*. The *G. multiradiatus* genome version used does not have chromosome 1c.

### Technical validation

Genomic DNA was evaluated using the Agilent bioanalyser for fragment size and smearing, Nanodrop for purity and Qubit for concentration. The sequencing data was subjected to quality assessment such as read quality and length distribution before assembly. Nanopore reads were assessed with NanoPlot v1.47.1^56^, a component of the NanoPack suite while the Omni-C and Illumina reads were assessed with fastQC v0.11.9^20^ and summarised with multiQC v1.35^21^ (Figure S2-S4). Alignment of the final Nanopore assembly to the original raw reads using Minimap2 v2.24^57^ with the option “-ax map-ont” resulted in an overall mapping rate of 99.75%. The Illumina reads used for polishing exhibited high mapping rates (99.74%) to the Nanopore assembly. The repeat annotation statistics were compared with the one reported for *Aphanius iberius* and other close relatives, which showed similar DNA repeat signature (Figure 5B). The RNA-seq reads from the older embryo samples used for annotation were fastP trimmed with average mapping rates of at least 89% to the reference genome using HISAT2 v2.2.1^39^ (Table S9). Gene assembly completeness prior and after annotation was assessed with BUSCO v5.8.3 Actinopterygii_odb10 and Cyprinodontiformes_odb10 datasets. Peptide completeness post annotation was assessed with Omark v0.4.1^43^ which indicated a high proportion of expected heirachical orthogroups (HOGs) within the cyprinodontoidei lineage of killifishes (Figure 6C, Table 2).

### Data Records

The raw sequencing data derived from Illumina, Nanopore and Omni-C sequencing were deposited in the NCBI SRA database with the accession number SRR34866668, SRR34866669, and SRR34866667, respectively (https://dataview.ncbi.nlm.nih.gov/object/PRJNA1295366?reviewer=2d1r6vth8s7l6mh0 <u>53lsvu0rhh</u>). The RNA-seq data used for annotation evidence were obtained from the ArrayExpress database under the accession number E-MTAB-16987^58^ (https://www.ebi.ac.uk/biostudies/arrayexpress/studies/E-MTAB-16987?key=e4c29bef-bb71-4238-a3db-f666d39ddf15). The resultant genome assembly was submitted to the NCBI Genome database with the accession number JBPVIJ000000000. The mitochondrial genome and its annotation were deposited in GenBank under the accession number PZ796930. Supplemental gene and repeat annotation files in different formats, plus other associated materials are available on Figshare^59^.

## Author contributions

R. Y. A., T.K., Y.S., M.R. and T. Kudoh conceived and designed the study. R.Y.A and R.M. performed animal tissue sampling, culling and storage. R.Y.A. isolated and prepared DNA samples for Nanopore and Illumina sequencing. P.O. and A.J. performed library construction and sequencing of the Illumina and Nanopore reads with preliminary QC. T.K., K.KN and Y.S. carried out all steps of the Omni-C sequencing and assembly polishing. R.Y.A., T.K., K.KN. and Y.S. carried out downstream bioinformatics analysis on the Illumina and Nanopore reads. R.Y.A conducted structural and functional annotation, prepared the initial manuscript draft and incorporated comments from all the other authors. All the authors read and approved the final version of the manuscript.

## Acknowledgements

R.Y.A., R.M., M.R. and T. Kudoh were funded by the NC3Rs (National Centre for the Replacement, Refinement and Reduction of Animals in Research) under the grant number NC/X001121/1.

## Code availability

No custom codes were developed for this study. All software were used according to the developers’ recommendations. The optional flags and specific parameters were used as described in the software documentation.

