## Supplemental_figures for "Chromosome-level assembly of the Arabian killifish as a novel biomedical model species"

**Table of Contents**

**Figure S1**. Phylogeny and sequence alignment of the cytochrome oxidase subunit-I (COI) genes from *Aphaniops dispar* and public Aphaniidae species.

**Figure S2**. Sequencing quality of the Illumina short reads used to polish the *Aphaniops dispar* genome.

**Figure S3**. Fragment distribution of the extracted high molecular weight (HMW) used for the Nanopore reads.

**Figure S4**. Sequencing quality of the Omni-c reads of the *Aphaniops dispar* genome.

**Figure S5**. Repeat element landscape in the entire *Aphanius dispar* genome and unplaced sequences.

**Figure S6**. Relative number of *A. dispar* specific orthogroups assigned to zebrafish orthologues using g:Profiler.

**Figure S7**. Top significantly enriched cellular component and molecular function GO terms within the *A. dispar* species specific orthologues.

List of supplementary tables.

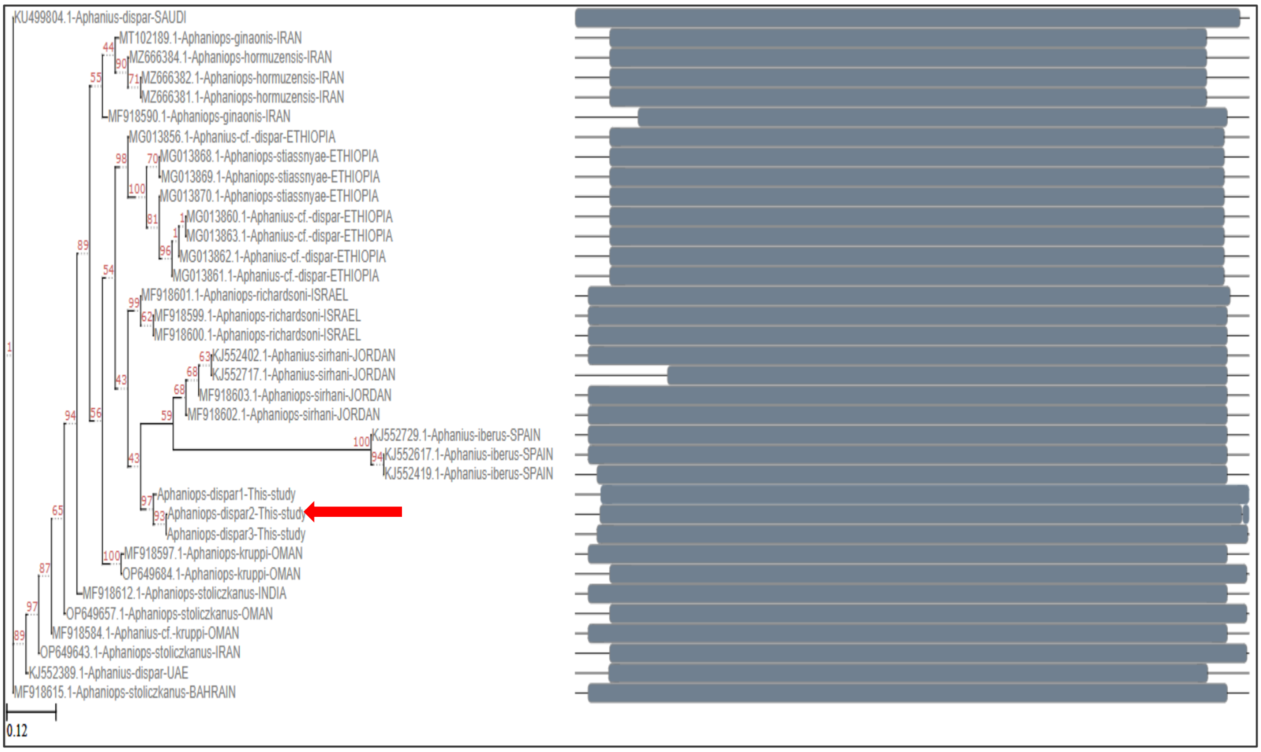


**Figure S1. Phylogeny and sequence alignment of the cytochrome oxidase subunit-I (COI) genes from *Aphaniops dispar* and public Aphaniidae species.** The red bar indicates the phylogenetic position of the three *Aphaniops dispar* samples metabarcoded for this study. The grey bars on the right are the COI sequence alignments displayed in aligned blocks from Mafft alignment output. Maximum-likelihood phylogenetic inference was performed using IQ-TREE with automatic model selection (ModelFinder) and 1,000 ultrafast bootstrap replicates. The node labels (in red) indicate bootstrap support values in percentages. Each sequence label represents the corresponding NCBI GenBank accession numbers, species name and original location in upper case. The scale bar indicates an average phylogenetic distance of 0.12 substitutions per site.

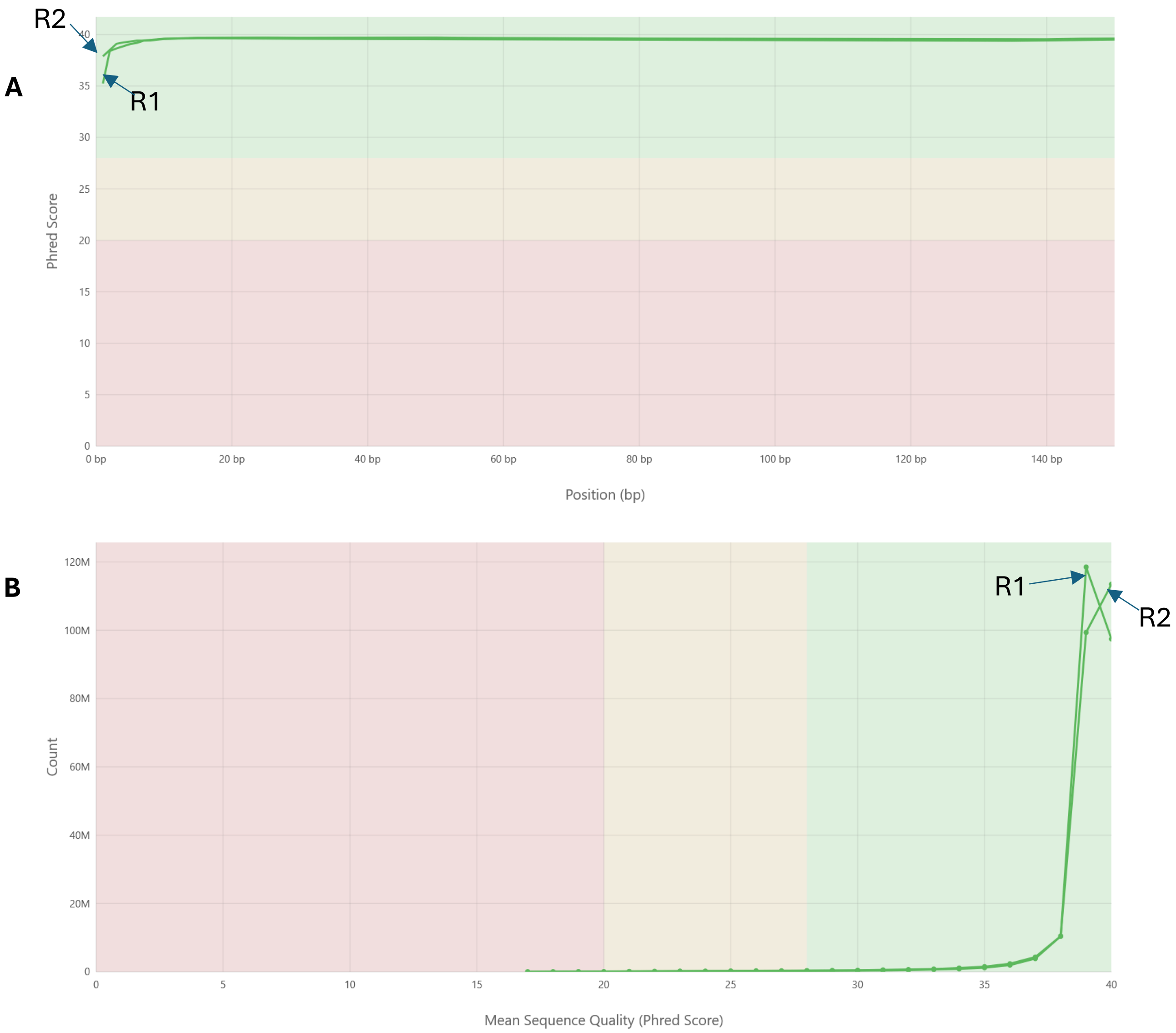


**Figure S2. Sequencing quality of the Illumina short reads used to polish the *Aphaniops dispar* genome.** Read 1 (R1) and Read 2 (R2) correspond to the forward reads and reverse reads of each paired-end fragment. A. Similar mean quality scores across the read lengths for forward and reverse reads. B. Per sequence quality scores showing sharp peaks between Q35-40 for both reads.

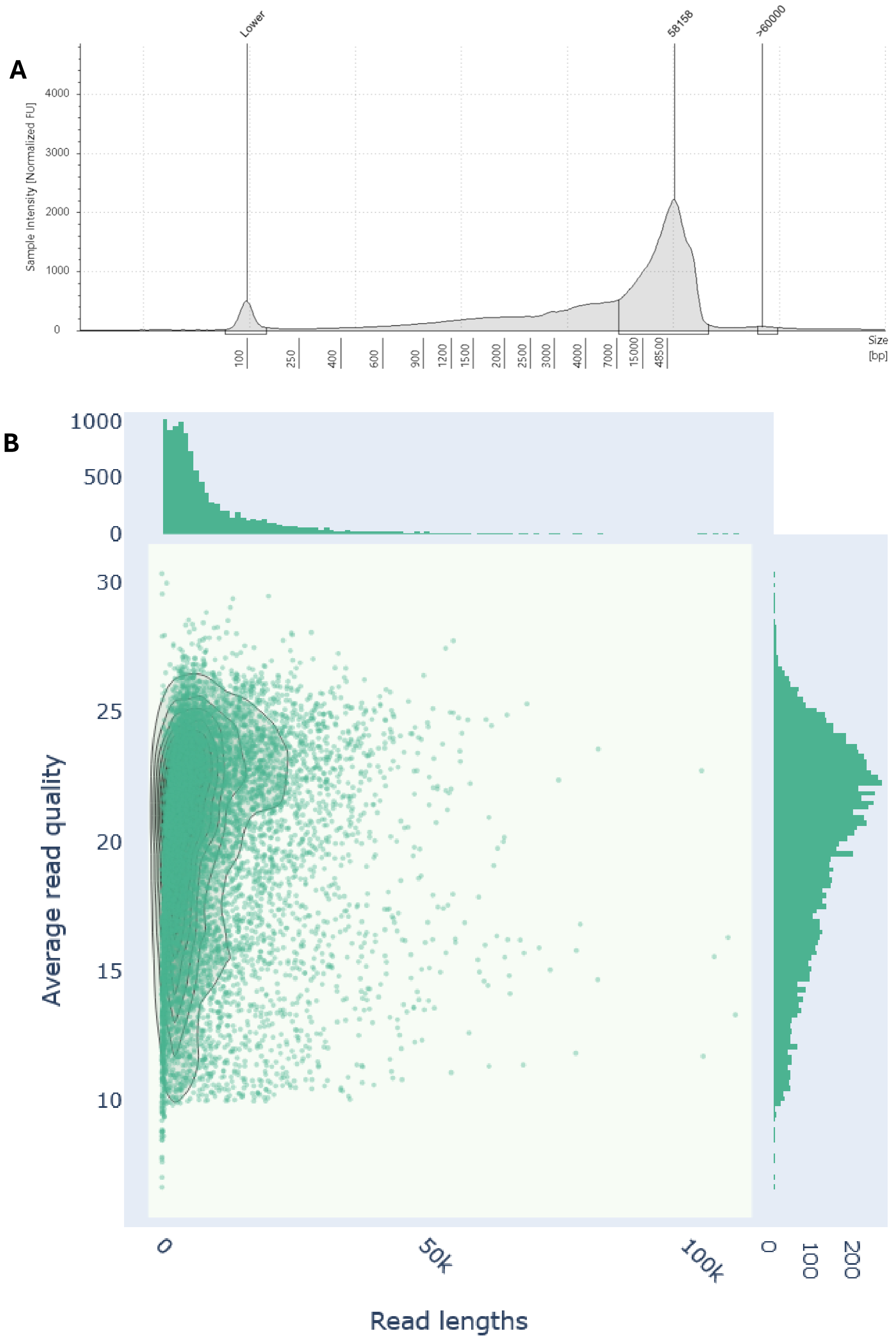


**Figure S3. Fragment distribution of the extracted high molecular weight (HMW) used for the Nanopore reads.** A. Size distribution of the extracted HMW DNA using the Agilent Fragment analyser. Traces of the small DNA fragments below 25Kb were eliminated with the Circulomics short read eliminator (SRE kit) prior sequencing. B. Read length and average read quality of the sequenced Nanopore long reads. The solid lines are kernel density estimates.

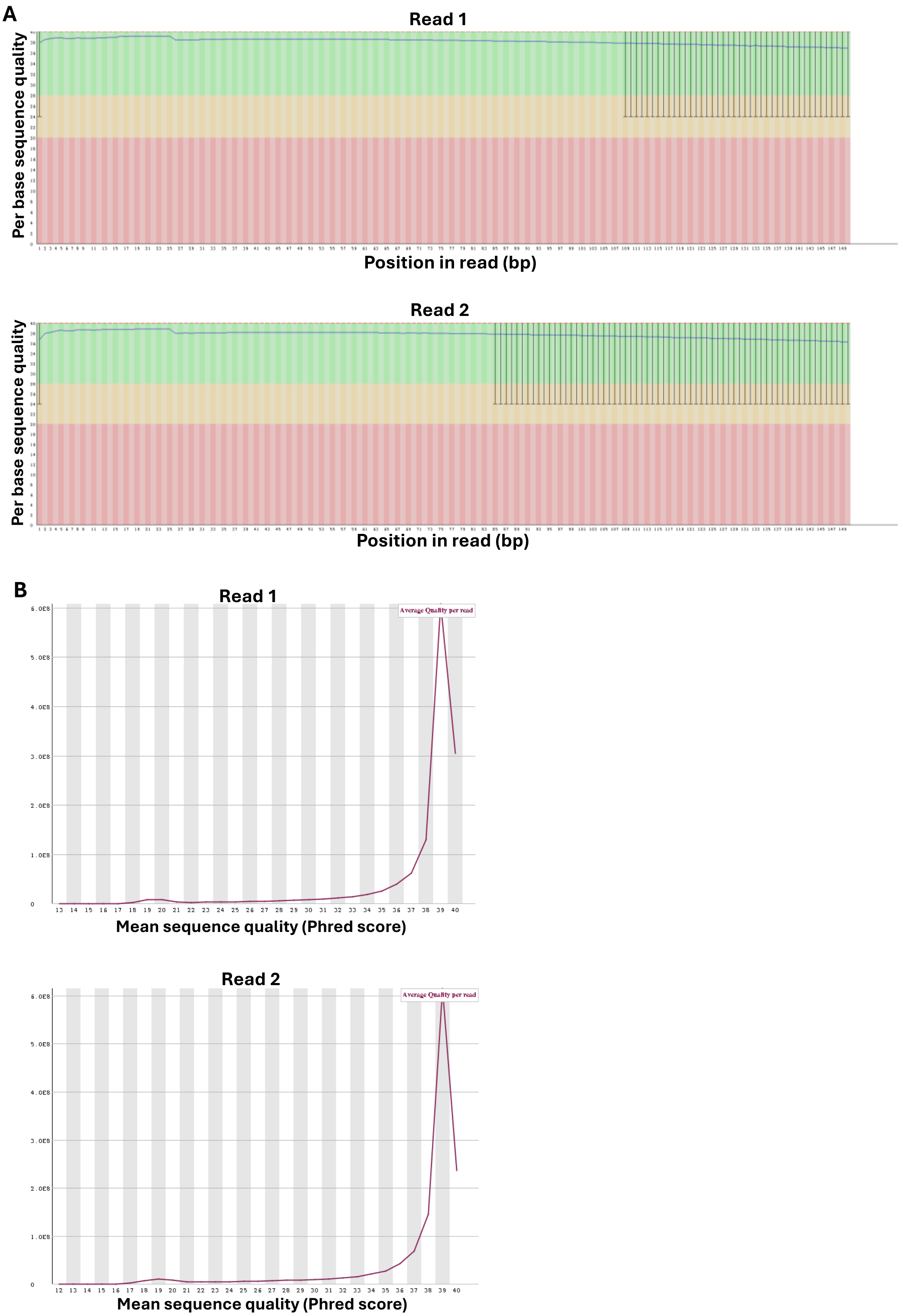


**Fig. S4. Sequencing quality of the Omni-c reads of the *Aphaniops dispar* genome.** (A) Per base sequence quality scores across all reads. (B) Per sequence quality scores.

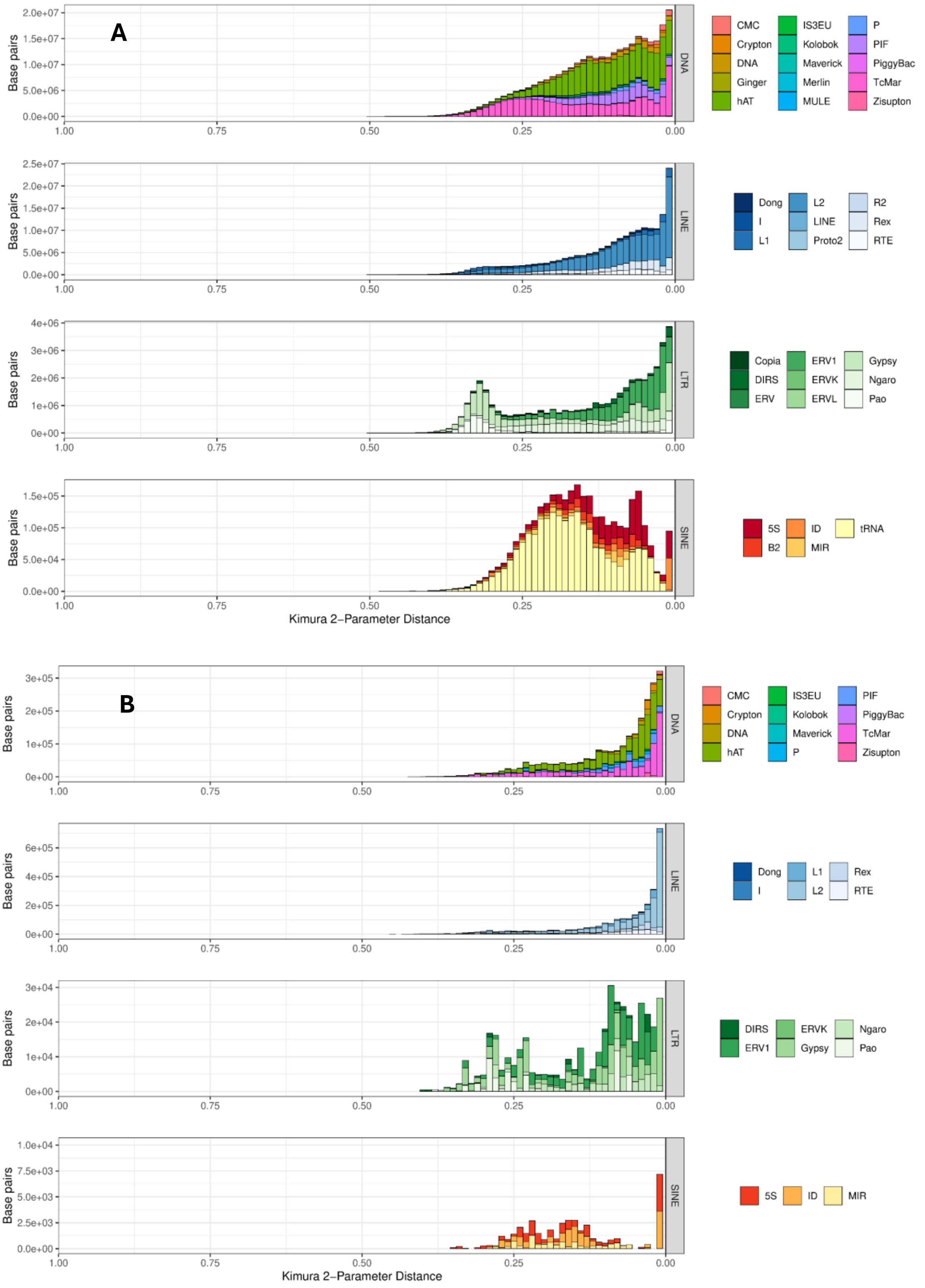


**Figure S5: Repeat element landscape in the entire *Aphanius dispar* genome and unplaced sequences.** A. Repeat landscape of the whole assembly (chromosome plus unplaced sequences). B. Repeat landscape of unplaced sequences. The x-axis represents the Kimura parameter distance, which estimates the evolutionary divergence of the younger repeat elements from their consensus sequences. The y-axis is the relative proportion of genome (in base pairs) occupied by each diverged repetitive element.

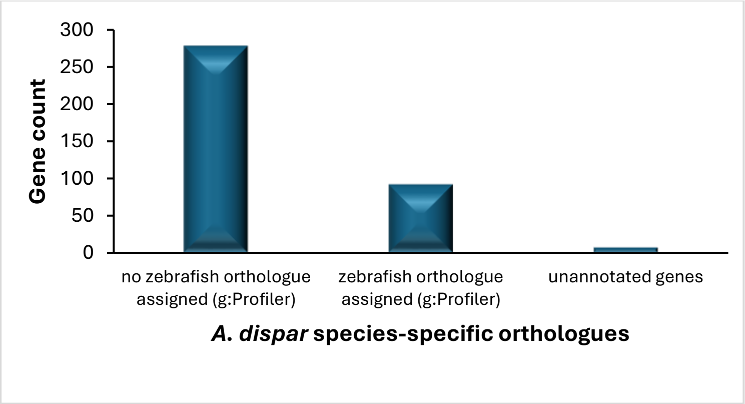


**Figure S6. Relative number of *A. dispar* specific orthogroups assigned to zebrafish orthologues using g:Profiler.**

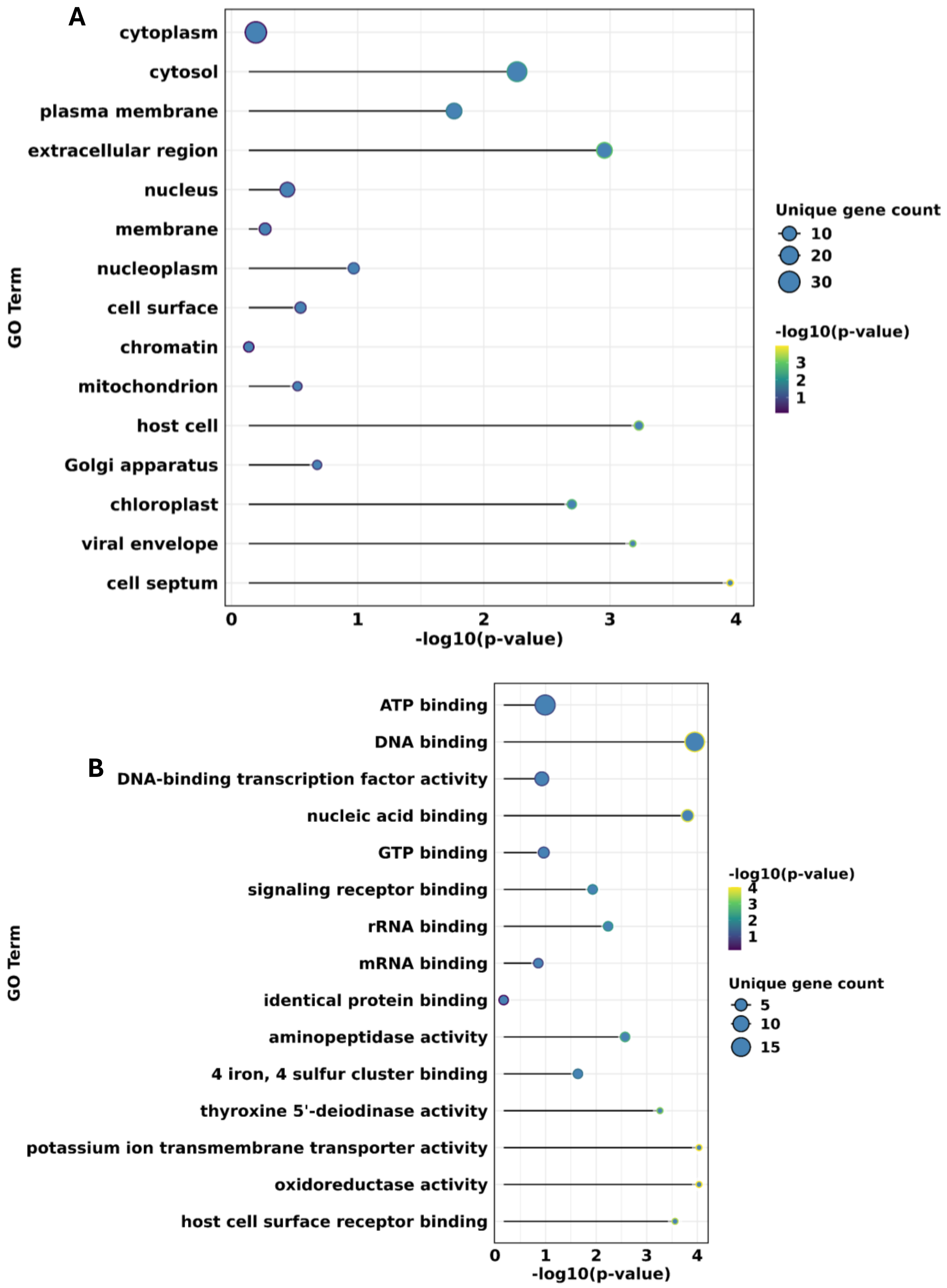


**Figure S7. Top significantly enriched cellular component and molecular function GO terms within the *A. dispar* species specific orthologues**. A: cellular component B: Molecular function.

**List of supplementary tables.**

**Table S1**. Metrics of the nanopore reads conducted with QUAST.
**Table S2**. Assembly statistics of the polished chromosome-scale assembly.
**Table S3**. *A. dispar* mitogenome statistics analysed with Quast.
**Table S4**. Structural variations between the *A.dispar* mitochondrial genome and that of its close relatives (*A. iberus* and *A. farsicus*).
**Table S5**. A summary of telomeric repeat statistics in all Chromosomes.
**Table S6**. Telomeric repeat identification in chromosomal and uplaced sequences with minimum repeat count of 100.
**Table S7**. Telomeric repeat identification in chromosomal and uplaced sequences with minimum repeat count of 25.
**Table S8**. Species used to generate protein evidence for *A. dispar* annotation.
**Table S9**. Alignment rates of embryonic transcriptome to the reference genome.
**Table S10**. A summary of AGAT statistics for genes and transcripts in the current BRAKER3 annotation round.
**Table S11**. List of missing BUSCO genes for Actinopterygii_0db10.
**Table S12**. BLASTP search results of BRAKER3-predicted proteins against proteins from 26 *Cyprinodontiformes* species in the RefSeq database.
**Table S13**. BLASTX search results of BRAKER3-predicted proteins against proteins from 26 *Cyprinodontiformes* species in the RefSeq database.
**Table S14**. BLASTP search results of BRAKER3-predicted proteins against the UNIPROT-SwissProt database.
**Table S15**. BLASTX search results of BRAKER3-predicted proteins against the UNIPROT-SwissProt database.
**Table S16**. BLASTX search results of BRAKER3-predicted nucleotides against zebrafish proteins in the RefSeq database.
**Table S17**. Search results of BRAKER3-predicted proteins against the eggNOG database.
**Table S18**. Functional annotation of the BRAKER3-predicted proteins using InterProScan.
**Table S19**. Chromosomal location, protein families and gene ontology of all annotated genes based on blastP searches in the UniProt_SwissProt database and the corresponding zebrafish hits.
**Table S20**. Species used for orthologous analysis and molecular phylogenetic analysis.
**Table S21**. Statistics per species from Orthofinder.
**Table S22**. Orthogroups from all species from Orthofinder.
**Table S23**. Orthogroups gene count from all species from Orthofinder.
**Table S24**. Gene ontology of *A. dispar* specific orthologues.
**Table S25**. Enriched GO terms within the *A. dispar* specific orthologues.
